## Supplementary Text for "Drug2ways: Reasoning over causal paths in biological networks for drug discovery"

### Additional File

#### Supplementary text

##### 1. Algorithm

Given the problem definition, we implemented two distinct functions, namely *all_paths* and *all_simple_paths* to count activatory and inhibitory paths considering any path (i.e. including *cyclic paths*) and considering *simple paths*, respectively.

Both algorithms make use of dynamic programming and memoization to save previous solutions between a pair of nodes and retrieve them every time they are needed, as part of another problem's solution. For *all_paths* any solution to a subproblem is enough to guarantee that we reach a valid solution to the final problem of counting paths between the source node and the target. However, in the case of *all_simple_paths*, the algorithm must be able to identify when a *cyclic path* is included in a stored solution and remove it from the total count. As partial solutions are stored as the number of accumulated paths between two nodes, there is no way to identify when a cycle may occur. One possible solution to this would be to store all paths found between two nodes and check which ones would form a cycle if added to the final solution. However, as discussed in **Subsection 4.1**, this is expensive in terms of compute time and memory, as the number of paths may grow exponentially and therefore, the space and time to store and check them does so as well. Alternatively, by keeping the list of intermediate nodes in an intermediate result, a potential cycle can be easily detected whenever a node $u$ is present in the list of intermediate nodes of one of its neighbors. Then, the node must be revisited to find and remove all paths with cycles. This, however, increases the complexity of the algorithm, as a node might be revisited more than once. We analyze in detail the scalability of both versions of the problem in **Subsection 2.4**. Following, we describe two algorithms which reason over all possible paths (**Algorithm 1**) and simple paths (**Algorithm 2**) and outline their pseudocode.

**Algorithm 1** describes the pseudocode of the main function to reason over all paths from node $u$ to $t$ with length less than or equal to $k$. We denote paths that activate a target as activations and paths that inhibit a target as inhibitions. These activations and inhibitions are counted separately. Line 2 and 3 are the stop conditions of the recursion. Line 2 checks whether node $u$ is equal to $t$, in which case, a path has been found. Line 3 checks if $k$ is 0, and if so, the maximum length has been reached without finding a path to the target node and the algorithm backtracks. Line 4 initializes a counter, a pair of integers which count activations and inhibitions from $u$ to $t$, respectively. Line 5 iterates over all nodes $v$ in the set of neighbors of $u$. Line 6 checks whether $v$ has been computed, and if so, the value is retrieved in Line 7. Otherwise, in Line 9 a recursive call is made to find all paths from $v$ to $t$ with a maximum length of $k - 1$. After obtaining the number of activations and inhibitions from *v* to *t*, the counter is updated accordingly in Lines 11 (if *u* activates *v*) and 13 (if *u* inhibits *v*). In Line 14, the result is saved in the cache.

---

**Algorithm 1**

Algorithm for counting paths with memoization and activation/inhibition counted separately.

---

**Require:**

$G = (V, E)$ and $u, v \in V$.

$k \in \mathbb{N}$.

**Ensure:**

$cache_{u \to t,k}$ contains the number of all paths from $u$ to $t$ of length at most $k$ that inhibit and activate $t$, separately.

Returns $cache_{u \to t,k}$.

1: **function** COUNT_PATHS$(G, u, t, k, cache)$

2: **if** $u = t$ **then return** $[1, 0]$

3: **else if** $k = 0$ **then return** $[0, 0]$

4: $paths_{u \to t,k} \leftarrow [0, 0]$

5: **for all** $v \in G.neighbors(u)$ **do**

6: **if** $paths_{v \to t,k}$ in $cache$ **then**

7: $paths_{v \to t,k} \leftarrow cache_{v \to t,k}$

8: **else**

9: $paths_{v \to t,k} \leftarrow$ COUNT_PATHS$(G, v, t, k-1, cache)$

10: **if** $(u, v) = 1$ **then** ▷ $u$ activates $v$

11: $paths_{u \to t,k} \leftarrow [\mathrm{paths}_{u \to t,k}[0] + paths_{v \to t,k}[0], paths_{u \to t,k}[1] + paths_{v \to t,k}[1]]$

12: **else** ▷ $u$ inhibits $v$

13: $paths_{u \to t,k} \leftarrow [\mathrm{paths}_{u \to t,k}[0] + paths_{v \to t,k}[1], paths_{u \to t,k}[1] + paths_{v \to t,k}[0]]$

14: $cache_{u \to t,k} \leftarrow paths_{u \to t,k}$

15: **return** $paths_{u \to t,k}$

---

**Algorithm 2** outlines the algorithm for reasoning over all simple paths with length less than or equal to *k*. Similar to **Algorithm 1**, given any pair of nodes *u, v* ∈ *V, s.t. (u, v)* ∈ *E* and any target node *t* ∈ *P,* activations and inhibitions from *u* to *t* are recursively obtained by first calculating activations and inhibitions from any *v* to *t*. However, in contrast to **Algorithm 1**, we keep track of the list of intermediate nodes to assist with the detection of cycles. Similar to **Algorithm 1,** Line 2 and 3 are the stop conditions of the recursion. Line 2 checks whether node *u* is equal to *t*, in which case, a path has been found. Line 3 checks whether *k* is 0, in which case, we have reached the maximum length without finding a path to the target node and we need to backtrack. Line 4 initializes the counter of paths for node *u* as an empty set. Line 5 iterates over all nodes *v* in the set of neighbors of *u*. Lines 6 and 7 check whether *v* was already computed and the value is retrieved. Otherwise, a recursive call is made in Line 9 to find all paths from *v* to *t* with a maximum length of *k - 1.* After obtaining the number of paths from $v$ to $t$, Line 10 determines if $u$ is any of the intermediate nodes of the paths from $u$ to $t$. In such a case, Line 11 revisits node $v$ to get all paths from $v$ to $t$ that include $u$ by calling the auxiliary function *get_paths_in_cycle*. Line 12 subtracts these paths from the total number of paths. Then, it is guaranteed that $paths_{v \to t,\, k-1}$ does not count any paths including $u$ and they can be added to $paths_{u \to t,\, k}$. This is done in Line 14, if $u$ activates $v$ and in Line 16 if $u$ inhibits $v$. Finally, $paths_{u \to t,\, k}$ is stored in cache and returned as the result of the function.

---

**Algorithm 2**

Algorithm for reasoning over simple paths. For this, cache needs to store number of paths through each intermediate node from $u$ to $t$ and activations/inhibitions are counted separately.

---

**Require:**

$G = (V, E)$ and $u,\, v \in V$.

$k \in \mathbb{N}$.

**Ensure:**

$cache_{u \to t,k}$ is a dictionary with all intermediate nodes present in all simple paths from $u$ to $t$ of length at most $k$ with the number of paths through each intermediate node. Inhibition and activation are counted separately. Returns $cache_{u \to t,k}$.

1: **function** COUNT_SIMPLE_PATHS$(G, u, t, k, cache)$
2:     **if** $u = t$ **then return** $\{t : [1, 0]\}$
3:     **else if** $k = 0$ **then return** $\varnothing$
4:     $paths_{u \to t,k} \leftarrow \varnothing$
5:     **for all** $v \in G.neighbors(u)$ **do**
6:         **if** $v \in cache$ **then**
7:             $paths_{v \to t,k-1}, \leftarrow cache_{v \to t,k-1}$
8:         **else**
9:             $paths_{v \to t,k-1} \leftarrow$ COUNT_SIMPLE_PATHS$(G, v, t, k-1, cache)$

10:         **if** $u \in paths_{v \to t,k-1}$ **then**   ▷ Node $u$ present in paths from $v \to t$
11:             $cycle\_paths \leftarrow$ GET_PATHS_IN_CYCLE$(G, v, t, u, k, cache)$
            ▷ Returns all paths from $cache_{v \to t,k-1}$ containing $u$.
12:             $paths_{v \to t,k-1} \leftarrow paths_{v \to t,k-1} - cycle\_paths$
            ▷ Remove paths that would form a cycle if considered.
13:         **if** $(u, v) = 1$ **then**   ▷ $u$ activates $v$
14:             $paths_{u \to t,k} \leftarrow [paths_{v \to t,k-1}[0], paths_{v \to t,k-1}[1]]$
15:         **else**   ▷ $u$ inhibits $v$
16:             $paths_{u \to t,k} \leftarrow [paths_{v \to t,k-1}[1], paths_{v \to t,k-1}[0]]$
17:     $cache[u] \leftarrow paths_{u \to t,k}$
18:     **return** $paths_{u \to t,k}$

---

### 2. Inferring causal interactions

The tables below describe the mappings between the original relationships as they appear in the networks and the two causal relationships (i.e., activation and inhibition) we subsequently infer.

| Relation | Equivalent Effect (Sign) |
|---|---|
| Increases | Activation (+1) |
| Regulates | Activation (+1) |
| Association (gene-disease) | Activation (+1) |
| Association (gene-phenotype) | Activation (+1) |
| Decreases | Inhibition (-1) |

**Supplementary Table 1. Relationships in the In-House network and their assigned polarity.** Mappings between original relations from source databases were made to equivalent, causal relationships (i.e., activation or inhibition). Gene-disease association edges were sourced from DisGeNet, each of which was the result of direct or indirect curation while OMIM's list of gene-disease associations provided links to genes. Thus, the confidence of gene-disease association edges from DisGeNet is 1. Because directionality was not provided for these gene-disease association relationships, they were inferred as activation edges. Similarly, gene-phenotype association edges were sourced from OpenBioLink and were inferred as activation edges.

| Relation | Equivalent Effect (Sign) |
|---|---|
| Drug-Activation-Gene | Activation (+1) |
| Gene-Activation-Gene | Activation (+1) |
| Gene-Phenotype | Activation (+1) |
| Gene-Disease | Activation (+1) |
| Drug-Binding activity-Gene | Activation (+1) |
| Drug-Inhibition-Gene | Inhibition (-1) |
| Gene-Inhibition-Gene | Inhibition (-1) |
| Drug-Binding inhibition-Gene | Inhibition (-1) |

**Supplementary Table 2. Relationships in OpenBioLink and their assigned polarity.** While original relations in OpenBioLink were directly translated to causal relations (i.e., activation and inhibition), gene-disease and gene-phenotype associations did not contain polarity and were inferred as activation edges.

### 3. Comparing distribution scores between the original and permuted networks

To assess the robustness of the predicted scores for the drug-disease pairs investigated in **Subsection 2.1**, we compared the distribution of the predicted scores in the same range of *lmax* for each of the two networks with the score distribution from permuted versions of the original networks, generated using XSWAP (Hanhijärvi et al., 2009) and the following implementation: https://github.com/dhimmel/xswap. We would like to mention that this algorithm maintains the original topology and only permutes the links between edges. Thus, source nodes (drugs) and target nodes (conditions) will maintain their original topology (no incoming and outgoing edges respectively). The results discussed in this section are attached in **Supplementary File 1.**

Before discussing the results, it is important to note the effect an increasing *lmax* can have on the network. Furthermore, for both versions (i.e., all paths and simple paths) a small *lmax* corresponds to a shorter and more direct path between the source (drug) and the target (disease). On the other hand, a larger *lmax* means that more paths can deviate towards distant parts of the network and the effect of feedforward loops can be captured to a greater extent by tracing various paths along them. Thus, due to this more extensive exploration, there is a point where a large *lmax* will result in paths that converge towards 0 (i.e., have an equal number of activation/inhibition interactions).

Across the two networks, we observe similar patterns in the distributions of the original versions, which contrast to those of the permuted versions, the latter of which also demonstrate marked similarities in their distribution trends. In OpenBioLink, we observe a similar pattern until *lmax*=8. This was in fact the reason why we chose this *lmax* as the upper bound for the validation. However, from this point the permuted network converges into a short-tailed distribution closely centered around 0 and the algorithm stops predicting candidate drug-disease pairs (i.e., there are no predictions close to +1 or -1). This pattern is even more pronounced for the In-House network. Here, while the original network converges and remains with its initial bimodal distribution (i.e., two consistent peaks on -1 and +1), the permuted version of the In-House network presents a short-tailed distribution that does not have any pairs in the tails of the distribution in *lmax* larger than 8. In summary, this experiment highlights that the original networks yield interesting hypotheses in contrast to the permuted networks which show no such biologically meaningful results.

### 4. Validation results for drugs predicted to activate a given indication

| **Network** | **All Paths** | | **Simple Paths** | |
|---|---|---|---|---|
| **-** | **7/7 Activate** | **6/7 Activate** | **7/7 Activate** | **6/7 Activate** |
| OpenBioLink | 0/0 (0%) | 1/3 **(33.33%)** | 0/0 (0%) | 4/9 **(44.44%)** |
| Permuted OpenBioLink | 0/0 (0%) | 0/0 (0%) | 0/0 (0%) | 0/0 (0%) |
| In-House | 11/18 **(61.11%)** | 56/494 **(11.34%)** | 9/11 **(81.82%)** | 56/488 **(11.48%)** |
| Permuted In-House | 0/0 (0%) | 0/6 (0%) | 0/0 (0%) | 0/7 (0%) |

**Supplementary Table 3. Results of the validation experiments focusing on prioritized drugs that activate an indication.** The table presents the validation experiments for each of the four networks (i..e, OpenBioLink, permuted OpenBioLink, In-House, and permuted In-House) using two versions of the algorithm (i.e., all paths and simple paths) based on two different criteria **(see Methods)**. For each experiment, we find a fewer number of true positive pairs prioritized in the top-ranked list that activate an indication, as compared to the number of pairs which inhibit **(Table 1)**.

| **Disease** | **Phenotypes** | **Source** |
|---|---|---|
| Cystic fibrosis of pancreas | Chronic obstructive pulmonary disease | HPO |
| | Exocrine pancreatic insufficiency | HPO |
| | Elevated sweat chloride | HPO |
| | Dehydration | HPO |
| | Chronic lung disease | HPO |
| | Meconium ileus | HPO |
| | Recurrent pneumonia | HPO |

**Supplementary Table 4. Phenotypes associated with cystic fibrosis of pancreas, the indication investigated in Subsection 2.2.**
