## Supplementary figures and images for "Drug2ways: Reasoning over causal paths in biological networks for drug discovery"

### all_paths_inhouse.png

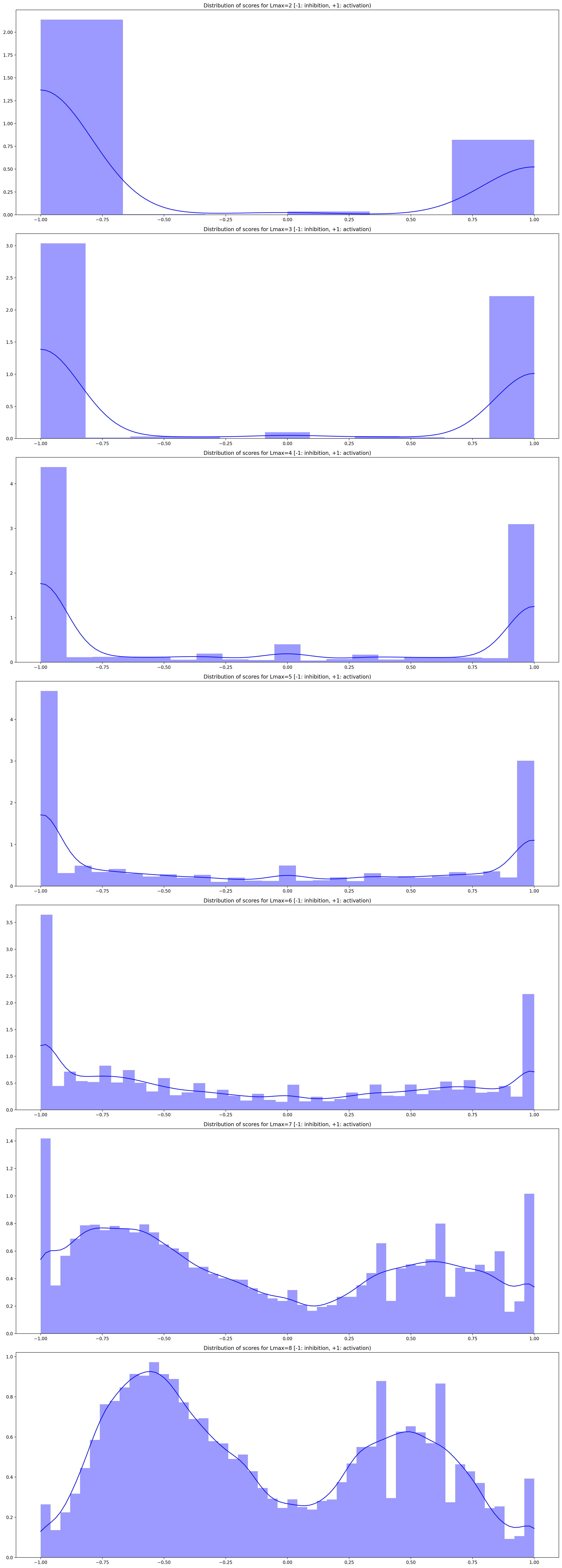

### all_paths_openbiolink.png

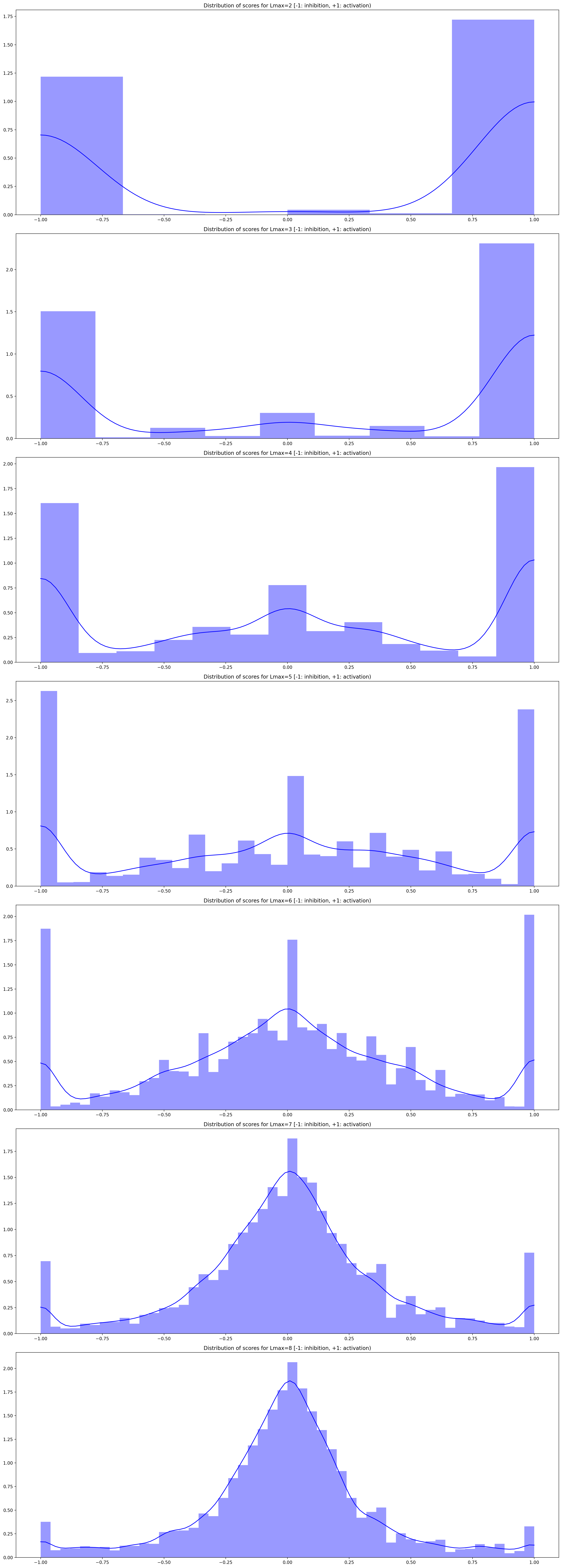

### permuted_ simple_paths_inhouse.png

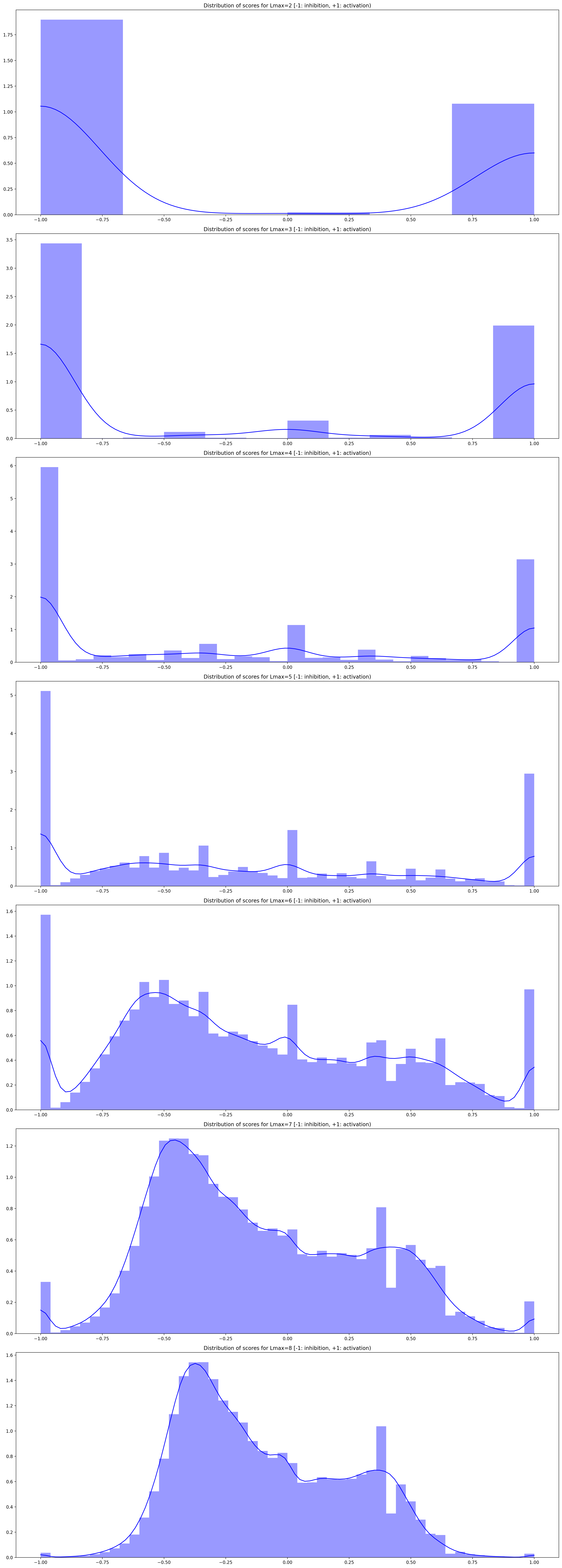

### permuted_all_paths_inhouse.png

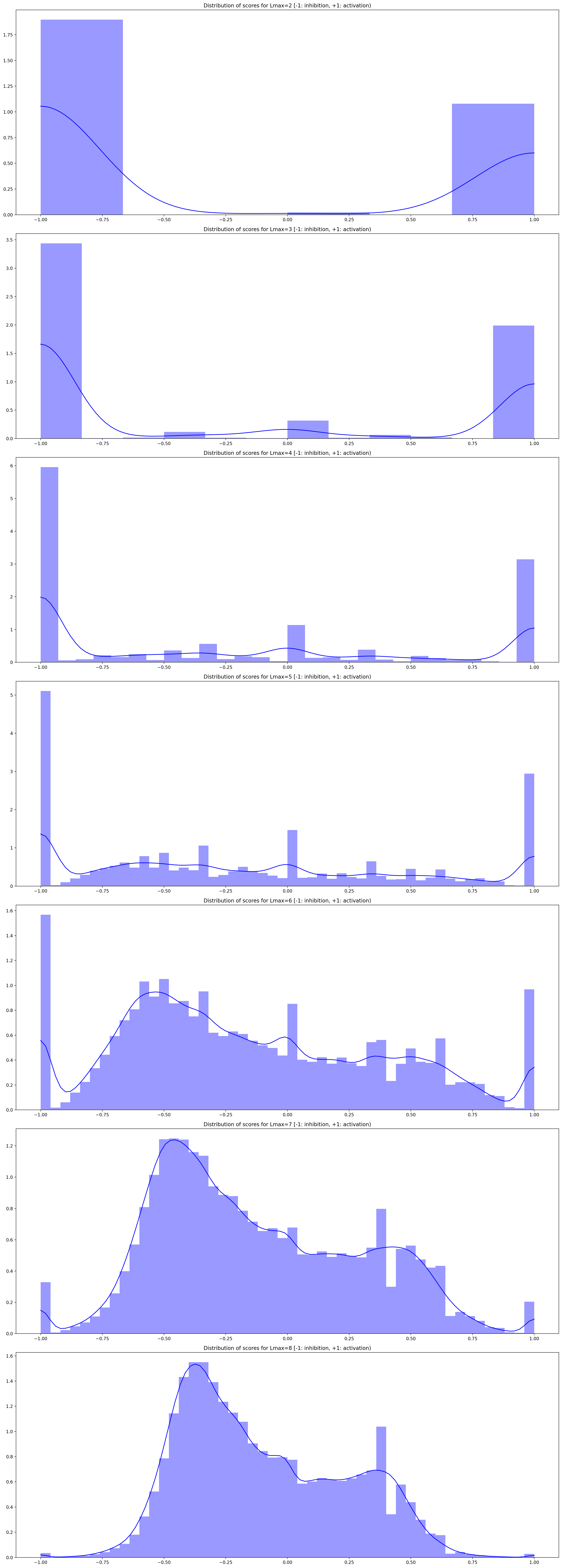

### permuted_all_paths_openbiolink.png

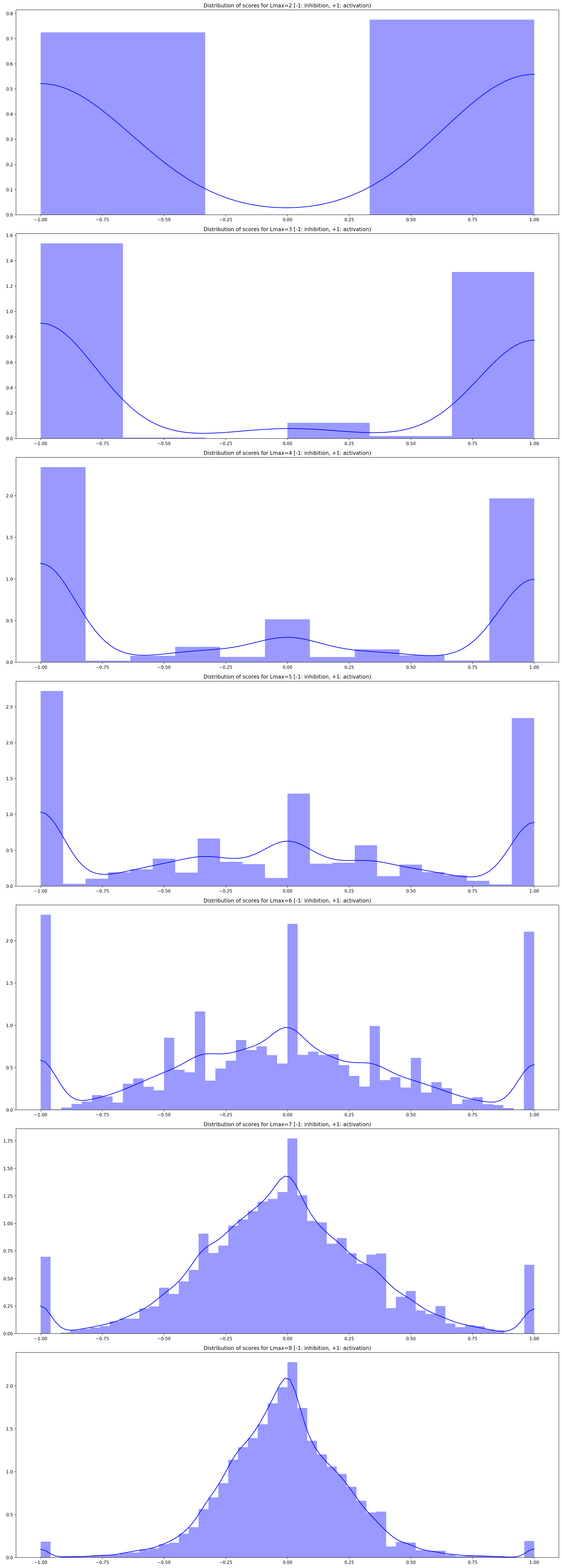

### permuted_simple_paths_openbiolink.png

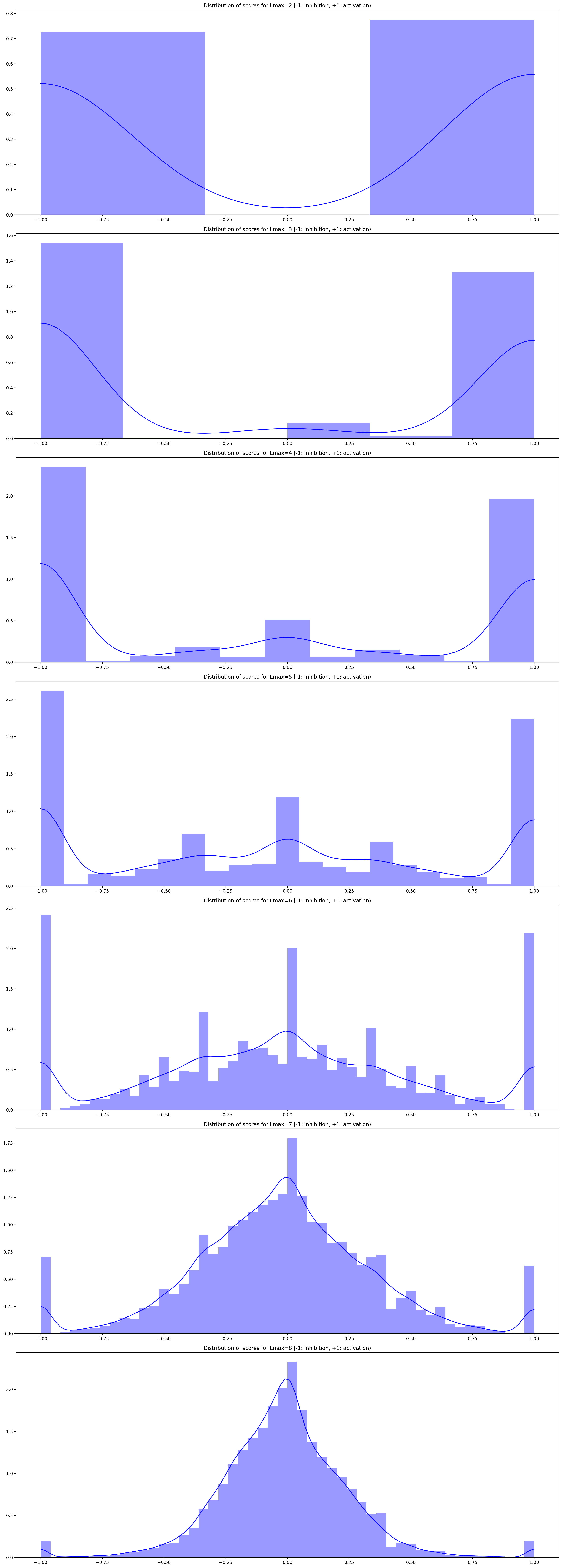

### simple_paths_inhouse.png

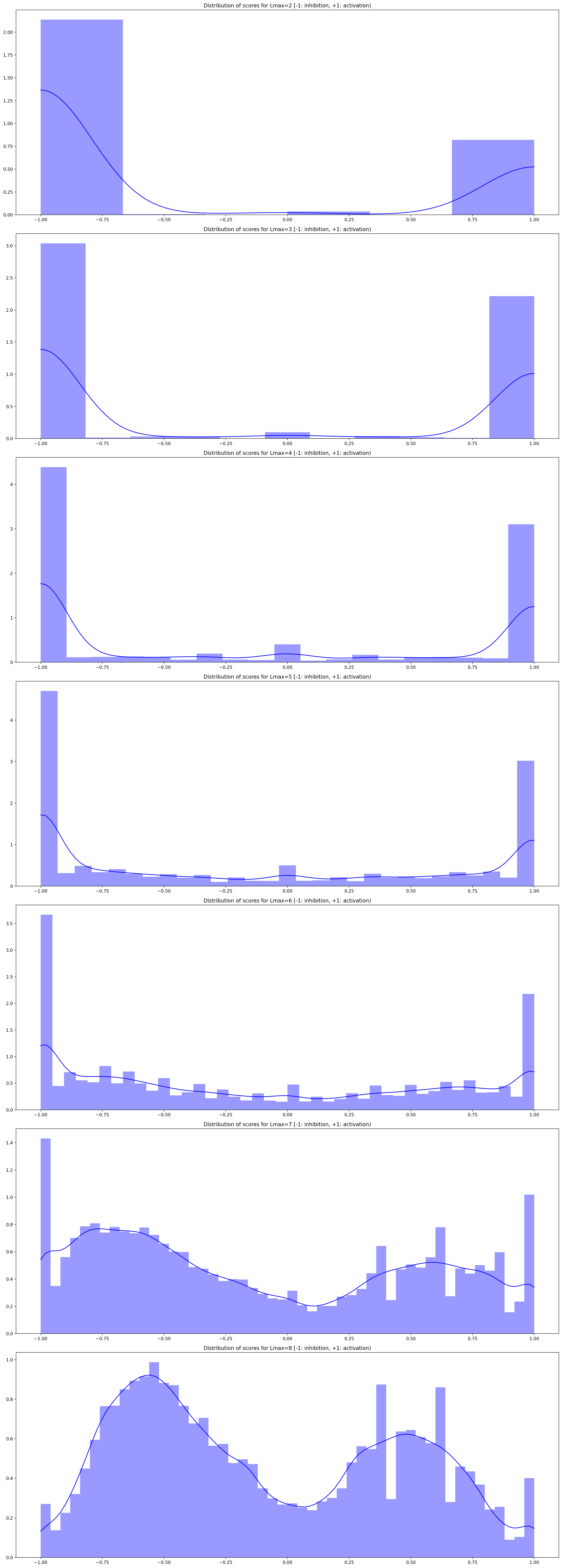

### simple_paths_openbiolink.png

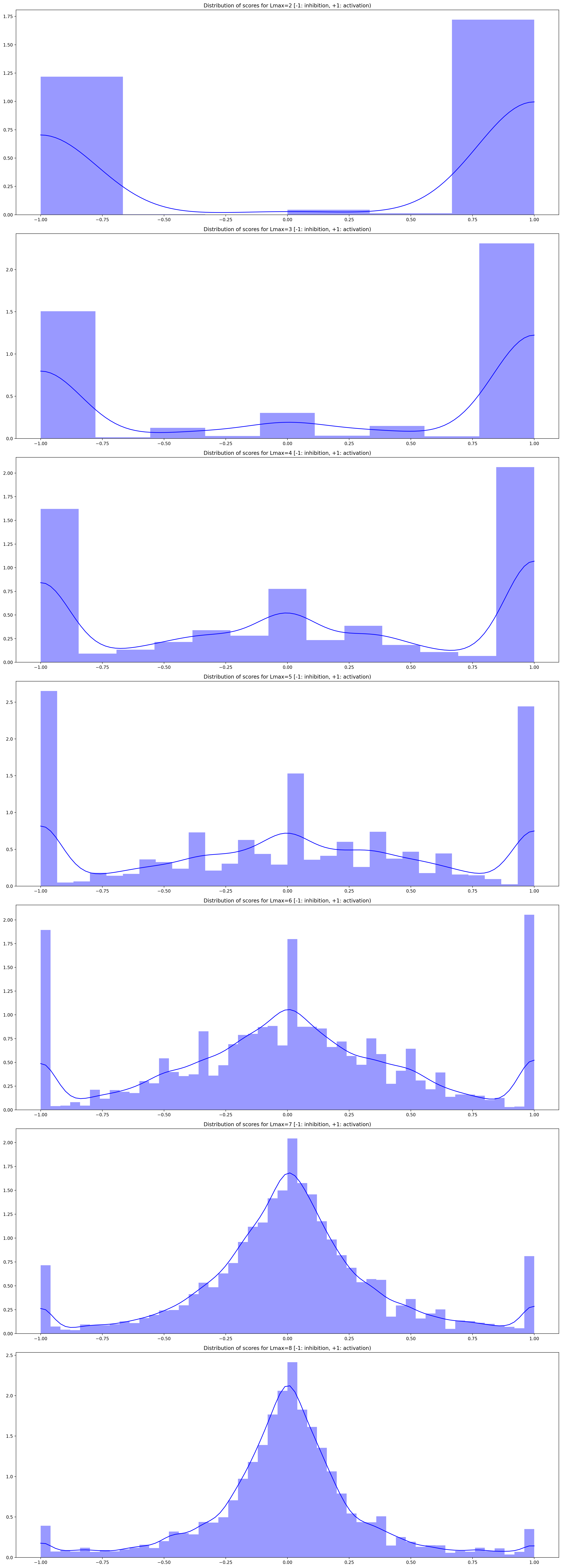
